# A Genome-engineered Bioartificial Implant for Autoregulated Anti-Cytokine Drug Delivery

**DOI:** 10.1101/535609

**Authors:** Yun-Rak Choi, Kelsey H. Collins, Luke E. Springer, Lara Pferdehirt, Alison K. Ross, Chia-Lung Wu, Franklin T. Moutos, Natalia S. Harasymowicz, Jonathan M. Brunger, Christine T.N. Pham, Farshid Guilak

## Abstract

Biologic drug therapies are effective treatments for autoimmune diseases such as rheumatoid arthritis (RA) but may cause significant adverse effects, as they are administered continuously at high doses that can suppress the immune system. Using CRISPR-Cas9 genome editing, we engineered stem cells containing a synthetic gene circuit expressing biologic drugs to antagonize interleukin-1 (IL-1) or tumor necrosis factor (TNF) in an autoregulated, feedback-controlled manner in response to activation of the endogenous chemokine (C-C) motif ligand 2 (Ccl2) promoter. To test this approach in vivo, cells were tissue-engineered into a stable cartilaginous construct and implanted subcutaneously in mice with inflammatory arthritis. Bioengineered anti-cytokine implants mitigated arthritis severity as measured by joint pain, structural damage, and systemic and local inflammation. The coupling of synthetic biology with tissue engineering promises a range of potential applications for treating chronic diseases using custom-designed cells that express therapeutic transgenes in response to dynamically changing biological signals.

---

Despite advances in the development of disease-modifying anti-rheumatic biologic drugs, approximately 40% of patients with rheumatoid arthritis (RA) fail to respond to treatment^1^. While the severity of RA fluctuates, biologic drugs are administered continuously at high concentrations, predisposing patients to significant adverse effects, such as increased risk of infection^2^. The development of therapeutics that can sense and respond to dynamically changing levels of endogenous inflammatory mediators may improve efficacy while mitigating the side effects of continuous biologic delivery^3-5^. Here, we used CRISPR-Cas9 genome engineering^6,7^ to create a self-regulating gene circuit in induced pluripotent stem cells (iPSCs). These cells were designed to produce anti-cytokine biologic drugs in response to inflammatory signals such as interleukin-1 (IL-1) or tumor necrosis factor *α* (TNF-*α*) by transcribing their biologic inhibitors in a feedback-controlled manner^3^, driven by the promoter of the *chemokine (C-C) motif ligand 2* (*Ccl2*) (Fig. 1)^8^. For delivery *in vivo*, iPSCs were differentiated into a cartilaginous implant and engineered into a cartilaginous implant that maintains the cells in a stable subcutaneous depot, allowing for sensing of systemic inflammation as well as free diffusion of the biologic drugs into the circulation. We demonstrated that these bioengineered implants provide dynamic, autoregulated delivery of anti-cytokine biologic drugs that mitigate structural damage, pain, and inflammation in a murine model of arthritis.

**Fig. 1.**
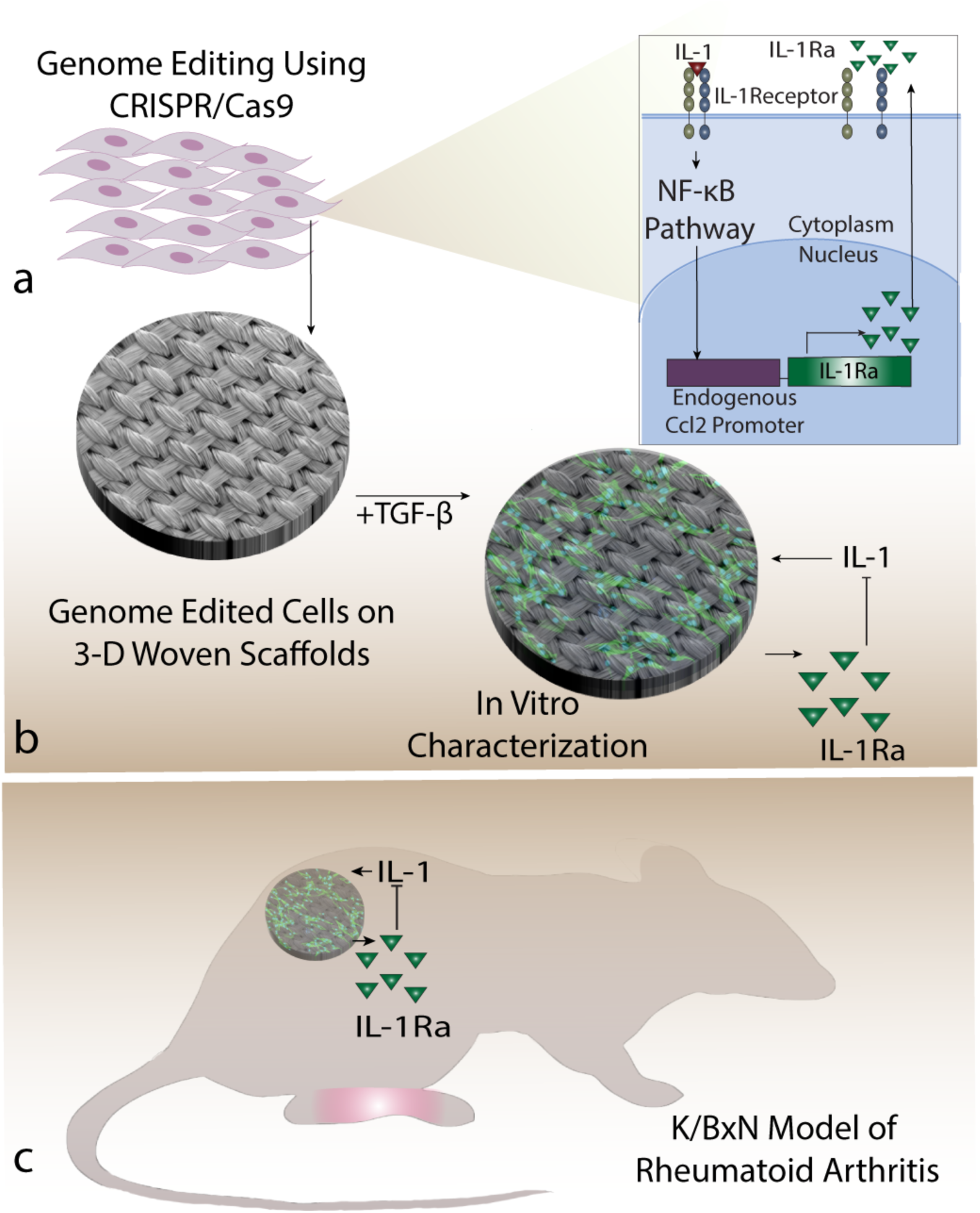
Scaffold-mediated Delivery of Genome-engineered Anti-Cytokine Cells. **a**. CRISPR-Cas9 genome editing was used to engineer stem cells containing a synthetic gene circuit that expresses the interleukin-1 receptor antagonist (IL-1Ra), an inhibitor of IL-1, in response to activation of the chemokine (C-C) motif ligand 2 (*Ccl2*) promoter. **b**. Cells were engineered to form stable cartilaginous constructs on 3D woven scaffolds in chondrogenic media. Constructs were tested *in vitro* to assess on-off dynamics in response to changing levels of IL-1. **c**. To test their efficacy, these anti-cytokine constructs were implanted subcutaneously in K/BxN model of rheumatoid arthritis.

Using CRISPR-Cas9 genome engineering, iPSCs were edited to insert either the gene for IL-1 receptor antagonist (*Il1rn*) or luciferase (*Luc*) at the *Ccl2* locus, creating a self-regulating gene circuit that transcribes *Il1rn* or *Luc* in response to inflammatory activation of *Ccl2* (referred as Ccl2-IL1Ra or Ccl2-Luc cells)^3^. We initially examined the therapeutic potential of Ccl2-Luc cells^9^ by intraperitoneal (IP) injection into a murine model of K/BxN serum transfer arthritis (STA)^10^. When triggered exogenously by a single IP injection of TNF-*α* to simulate an RA “flare”, we observed robust luciferase expression (Suppl. Fig. 1a) that recovered after the first 24 h (Suppl. Fig. 1b). In the context of K/BxN STA, Ccl2-IL1Ra cells delivered by IP injection did not mitigate clinical scores (p=0.10, Suppl. Fig. 1c), but significantly suppressed ankle swelling (13.8% reduction on day 6) and mitigated pain sensitivity (25% reduction in untreated animals, no change in IL-1Ra treated animals; Suppl. Fig. 1d-f).

Due to the lack of long-term cell engraftment by IP injection (as measured by luciferase expression after 24h), as well as the modest mitigation of disease severity, we used an alternative tissue-engineering approach to create a cartilaginous construct to provide stable engraftment. Cells were seeded onto 3D woven poly(ε-caprolactone) (PCL) scaffolds (Fig. 2a)^11^ and chondrogenically differentiated^12^ over 21 days into chondrocyte-like cells that produced a proteoglycan-rich matrix (Fig. 2b-d). In culture, the Ccl2-IL1Ra bioartificial implants sensed and responded to stimulation with IL-1*α* by producing IL-1Ra (Fig. 2e) in a feedback-controlled manner (Fig. 2f,g). A second set of cells were similarly engineered to produce soluble TNF receptor 1 (sTNFR1), and implants created from Ccl2-sTNFR1 cells behaved similarly *in vitro* (Suppl. Fig. 2). The cell-secreted anti-cytokine biologics suppressed the response to IL-1*α* or TNF-*α*, reducing mRNA levels for proinflammatory mediators *Il6* and *Ccl2*, and for matrix metalloproteinases, *Mmp9* and *Mmp13* (Fig. 2h-k, Suppl. Fig. 2), as compared to the control Ccl2-Luc construct.

**Fig. 2.**
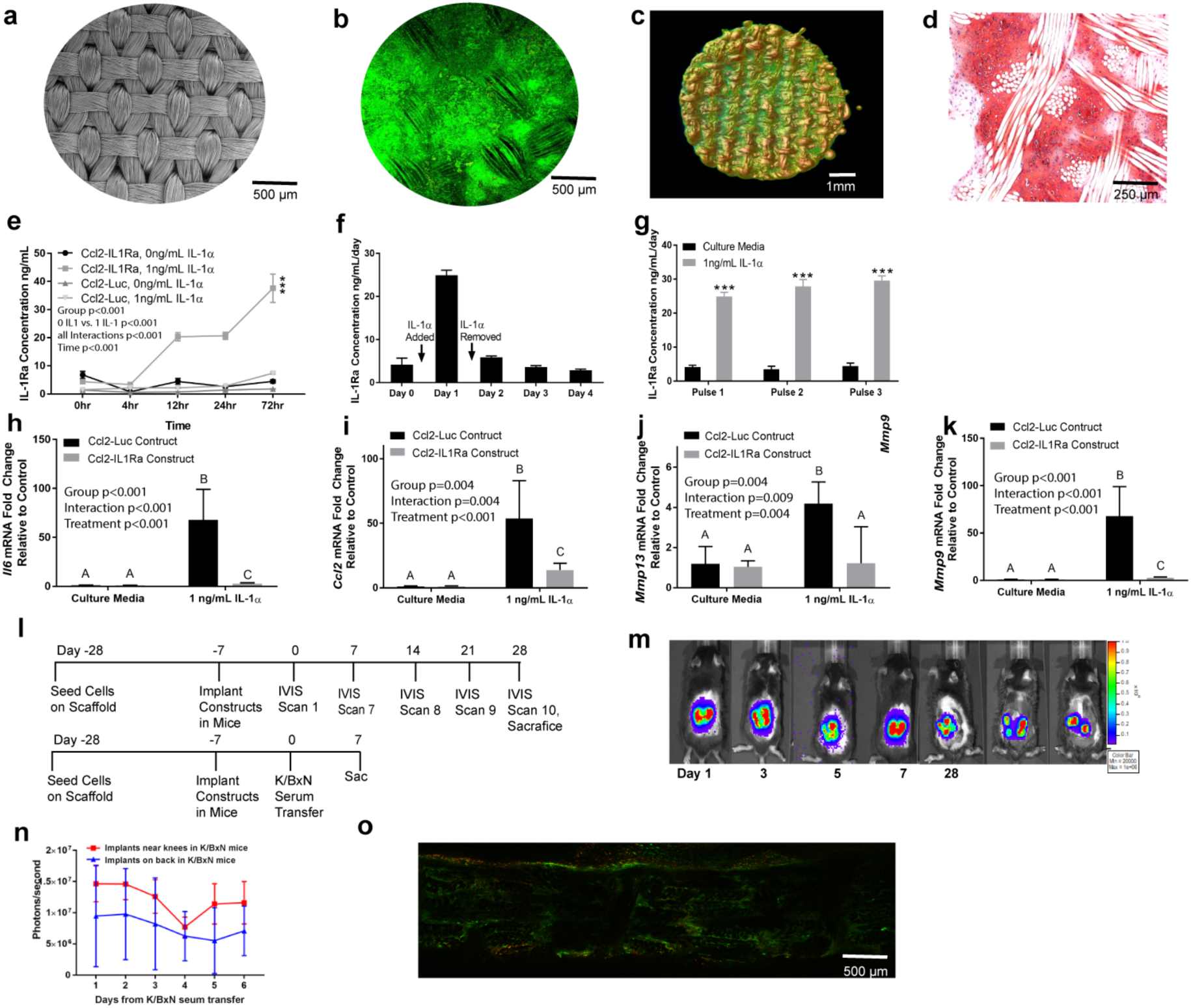
Tissue-engineered Bioartificial Implants Sense and Respond to Inflammation in a Feedback-controlled Manner. **a-d**. Genome-engineered stem cells on porous 3D woven scaffolds (**a**, electron microscopy) were differentiated into chondrocyte-like cells that infiltrated the scaffold and filled the scaffold pores with cartilaginous matrix as shown by **b**. confocal microscopy; **c**. nanoCT; **d**. Safranin O staining. **e**. Ccl2-IL1Ra constructs produced IL-1Ra in response to IL-1*α* and **f**. decreased IL-1Ra production rapidly after IL-1*α* withdrawal. **g**. This response was repeatable over multiple cycles of iterative stimulation (n = 5 per group). **h-k**. Ccl2-IL-1Ra constructs (n = 6) mitigated mRNA levels for pro-inflammatory mediators as compared to Ccl2-Luc constructs (n = 5). **l-o**. When implanted in K/BxN model of rheumatoid arthritis, Ccl2-Luc implants expressed consistent luciferase (**m**,**n**) and demonstrated high cell viability (**o**) upon explantation for confocal microscopy (n ≥ 3 per group). All values represent mean ± SEM. In **e**, three-way repeated measured ANOVA, in **g**, Students’ t-test: *** indicates p < 0.001. In **h-k** and **n**, two-way ANOVA: different letters indicate p < 0.001 by Tukey’s post-hoc test.

These bioartificial constructs were implanted subcutaneously onto the dorsal aspect of K/BxN transgenic F1 mice, which develop spontaneous arthritis, as well as mice with K/BxN STA^10^. Ccl2-Luc constructs exhibited high luminescence relative to background levels, indicating responsiveness of the constructs *in vivo*, as well as cell survival in a chronic inflammatory environment up to 5 weeks after implantation in F1 K/BxN mice (Fig. 2l,m). Ccl2-Luc constructs implanted in the K/BxN STA model exhibited similar luminescence signals for 7 days when they were implanted subcutaneously near the knees or on the back (Fig. 2n). Explanted Ccl2-Luc constructs from F1 K/BxN mice at 5 weeks also confirmed high cell viability via live/dead staining by confocal microscopy (Fig. 2o).

Chondrocyte-like cells in a cartilage matrix exhibit minimal migration and require no vasculature for long-term survival, relying on diffusion for nutrient transport^13-16^. To determine the ability of this self-regulating implant to mitigate RA disease severity, constructs were implanted on the dorsal aspect of C57BL/6 mice and allowed to recover for 7 days followed by intravenous K/BxN serum administration on day 0 (Fig. 3a). Animals receiving Ccl2-IL1Ra implants demonstrated significant improvements in RA severity, including 40% reductions in both clinical scores and ankle thickness, compared to control animals that received Ccl2-Luc constructs or no implant (Fig. 3b,c). Histologic analysis showed that animals with Ccl2-IL1Ra implants demonstrated significantly lower inflammation scores and had reduced cartilage degradation and proteoglycan loss (Fig. 3 d-f) when compared to animals with Ccl2-Luc implants. Mice receiving Ccl2-IL1Ra constructs also exhibited significantly higher serum levels of IL-1Ra when compared to controls (Fig. 3g,h). Serum IL-1Ra concentrations correlated with clinical scores and ankle thickness (Fig. 3i,j). The reduction in disease severity in mice with Ccl2-IL1Ra constructs was accompanied by a significant decrease in mechanical pain sensitivity as measured by algometry and Electronic Von Frey pain tests, whereas the control groups exhibited increased pain sensitivity (reduced pain thresholds) (Fig. 3k,l). The pain threshold of each animal also significantly correlated with serum levels of IL-1Ra (Fig. 3m,n). Neither Ccl2-sTNFR1 (Suppl. Fig. 3) nor reducing the dose of Ccl2-IL1Ra constructs by half (Suppl. Fig. 4) significantly mitigated structural damage from K/BxN STA, although both approaches significantly mitigated an increase in pain sensitivity (Suppl. Fig. 3, 4). These data corroborate previous reports that TNF-α and IL-1α signaling are both directly and indirectly associated with the onset of pain in K/BxN STA^17^.

**Fig. 3.**
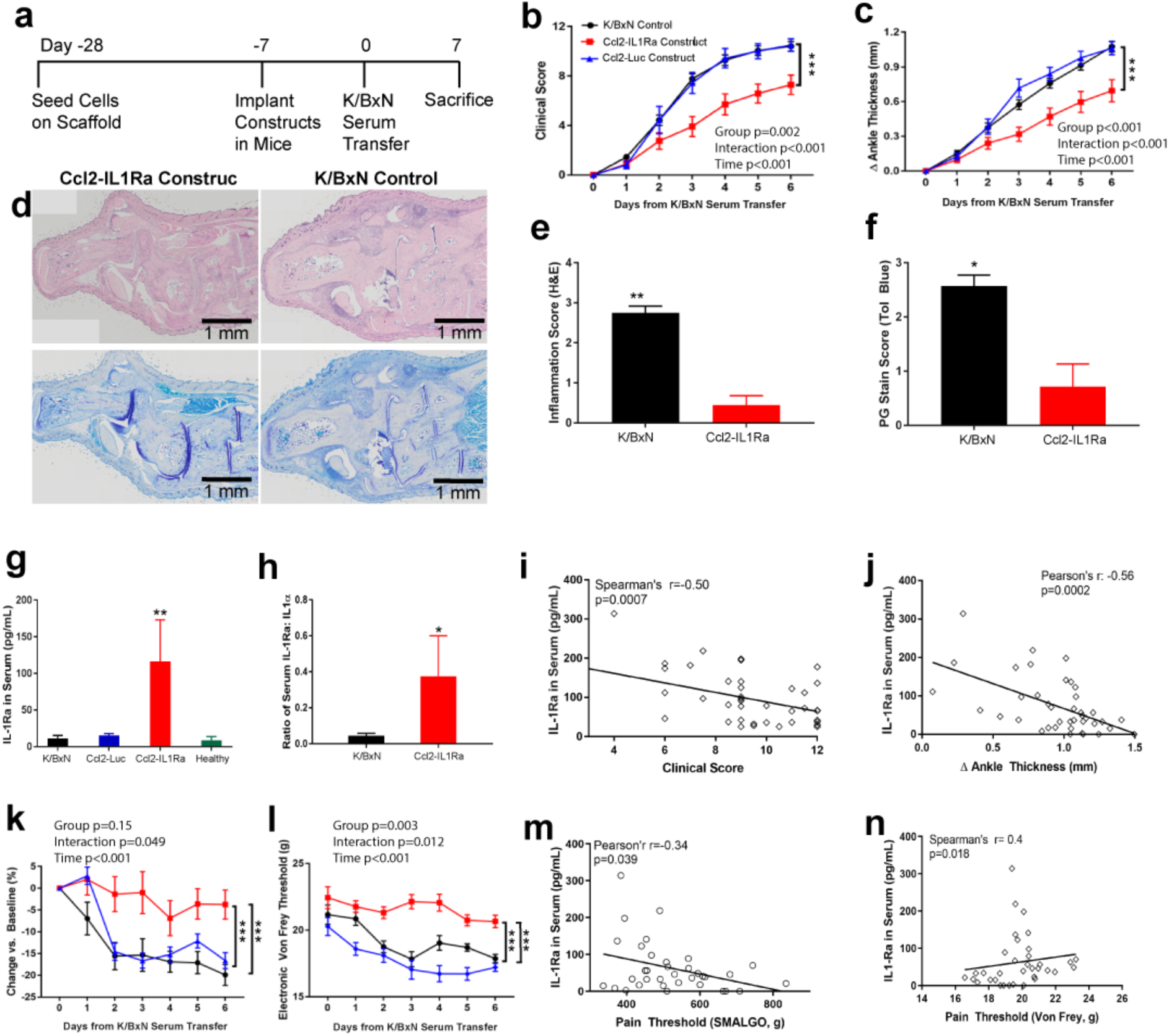
Ccl2-IL1Ra Bioartificial Implants Mitigate Disease Severity, Inflammation, and Pain in Mice with K/BxN Serum Transfer Arthritis. **a-c**, Ccl2-IL1Ra implants mitigated disease severity as measure by clinical scores and ankle thickness, in contrast to control groups (n = 20, 12, 8, respectively). **d-f**. At sacrifice, these implants reduced inflammation (**e**, n = 8, 9, left to right) and maintained cartilage integrity (**f**. n = 7 per group). **g-j**. IL-1Ra levels and ratio of IL-1:IL-1Ra in serum were increased in mice with Ccl2-IL1Ra implants compared to all other groups and IL-1Ra concentration in serum had negative relations with clinical scores, ankle thickness (**g**. n = 10, 6, 12, 8, left to right; **h**. n= 10 per group). **k-n**. For pain measurements, mice receiving Ccl2-IL1Ra implants maintained their pain thresholds, indicated by algometry and Electronic Von Frey pain tests in contrast to all other groups and significantly related with IL-1Ra concentration in serum (n = 20, 12, 8, respectively). All values represent mean ± SEM. In **b, c, k**, and **l**, two-way repeated measures ANOVA; in **e** and **f**, one-sided Mann-Whitney *U* test; in **g**, one-way ANOVA; in **h**, Students’ t-test. *P < 0.05, **P < 0.01, ***P < 0.001.

**Fig. 4.**
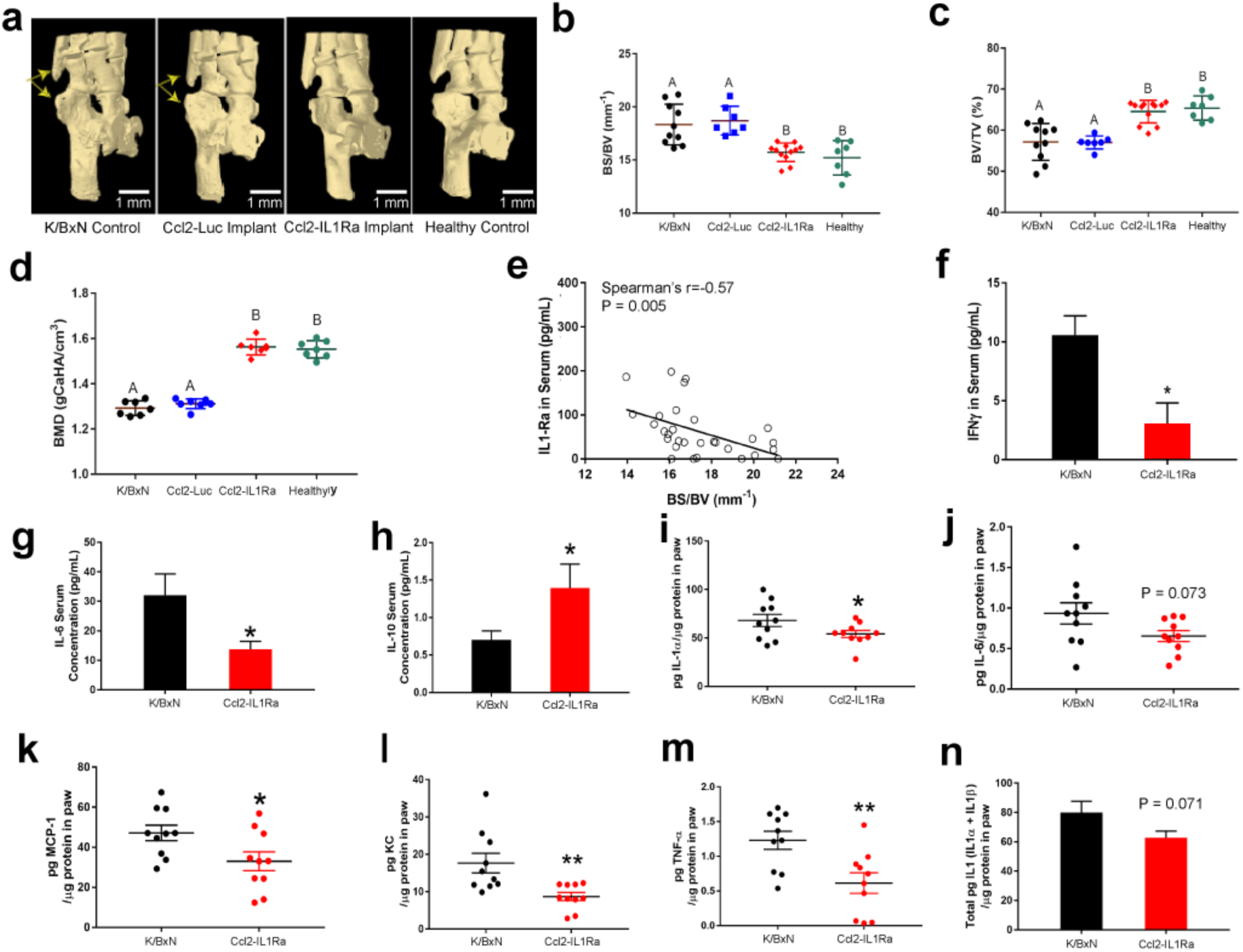
Ccl2-IL1Ra Bioartificial Implants Prevent Bone Damage and Mitigate Endogenous Inflammation in Mice with K/BxN Serum Transfer Arthritis. **a-e**. Mice treated with Ccl2-IL1Ra implants demonstrated reduced bone damage by microCT as quantified by **b**. the ratio of bone surface to bone volume (BS/BV); **c**. bone volume per total volume (BV/TV); **d**. bone mineral density (BMD); (n = 10, 7, 12, 7, left to right). **e**. Significant relationships were observed between BS/BV and serum levels for IL-1Ra. **f-n**. Mice receiving Ccl2-IL1Ra implants also showed less systemic (**f-h**) and local inflammation (**i-n**) as compared to mice with Ccl2-Luc implants or no implants (n = 10 per group). All values represent the mean fold change in expression ± SEM. In **b-d**, one-way ANOVA; in **f-n**, Students’ t-test; *P < 0.05, ** P < 0.01, different letters indicate P < 0.001 by Tukey’s post-hoc test.

Microcomputed tomography (microCT) of bone structure showed that mice receiving Ccl2-IL1Ra implants had minimal or no bone erosion on day 7 following K/BxN serum transfer, whereas mice with Ccl2-Luc implants or no implant exhibited highly erosive disease as demonstrated by a higher ratio of bone surface to bone volume (BS/BV) and a lower ratio of bone volume per total volume (BV/TV) (Fig. 4a-c). Histologic analysis was also concordant with the microCT observations for bone erosions (Fig. 3d). Ccl2-IL1Ra implants also showed protective effects on bone mineral density (BMD) (Fig. 4d). Quantitative measures of bony erosions (BS/BV) were negatively correlated with serum levels of IL-1Ra (Fig. 4e). Bony erosions are frequently observed in RA^18^ and are associated with disease severity and poor functional outcome. However, most biologics currently clinically available rarely inhibit both bone erosions and inflammation^18^. The ability to mitigate both inflammation and structural damage using this approach suggests that inhibition of IL-1 signaling in an autoregulated manner may provide added therapeutic advantage in the prevention of bone loss during inflammatory arthritis.

Mice receiving Ccl2-IL1Ra implants showed significant reductions in their inflammatory cytokine profiles as compared to control animals. Using a multiplexed cytokine assay, we observed significant reductions of serum levels for IFN-γ and IL-6, and an increase in serum levels for IL-10 in treated animals (Fig. 4f-h). In the paw lysate of treated animals, we observed significant reductions in levels for IL-1α, IL-6, MCP-1, KC, and TNF-*α* in comparison to control animals (Fig. 4i-m). There was a trend toward significance in the total level of IL-1 (IL-1α + IL-1β) in animals that received Ccl2-IL1Ra implants compared to controls (p=0.07, Fig. 4n).

In summary, we developed a genome-engineered implantable drug delivery system that can automatically sense and respond to inflammatory cytokines to produce therapeutic levels of anti-cytokine biologics in an auto-regulated manner. By engineering iPSCs to form a cartilaginous matrix, the implants exhibited long-term viability and function *in vivo*, with the ability to serve as a continuous source for biologic drugs. This platform represents a clinically-relevant and translational delivery strategy that may be applicable to a wide variety of diseases for long-term therapeutic control. While protein-based drug delivery, including enzyme-activated systems, have been successful in a number of applications^19-22^, such approaches are generally based on biodegradable materials and cannot easily supply continuously biologic molecules for long-term delivery, unlike the rapid dynamic stimulus-response functions that living cells can provide. In previous studies, “designer” or “smart” cell approaches have been used to develop cell therapies for metabolic diseases such as diabetes, but have been based on transient transfection or viral gene delivery, resulting in unpredictable gene insertion sites or copy numbers^5^. This study is among the first to demonstrate use of CRISPR-Cas9 gene editing for precise control of the stimulus-response locus *in vivo*^3^, as well as the potential for tunability and genome editing of cytokine receptors for treating model systems of an inflammatory condition^23,24^. Applying tissue engineering in combination with synthetic biology^25^ to develop artificial gene circuits that can sense and respond dynamically to disease markers may provide new opportunities for developing safe and effective therapies for chronic autoimmune diseases such as RA^26^.

## Methods

### Cell Culture and In Vitro Assays

Murine induced pluripotent stem cells (iPSCs) were engineered to incorporate a synthetic gene circuit to drive the expression of either *Il1rn* or firefly luciferase (*Luc*) via the *Ccl2* locus^3^. These iPSCs were predifferentiated in micromass with BMP-4^12^, seeded onto 3D woven poly(ε-caprolactone) (PCL) scaffolds (1e6 cells/scaffold), and cultured for 21 days in serum-free chondrogenic medium containing Dulbecco’s Modified Eagle Medium - High Glucose (DMEM-HG), non-essential amino acids, 2-mercaptoethanol, ITS+ premix (BD), penicillin-streptomycin (Gibco), 50 μg/mL L-ascorbic acid 2-phosphate, 40 μg/mL L-proline, 10 ng/mL transforming growth factor beta 3 (TGF-β3, R&D Systems) and 100nM dexamethasone^12,27^ to make implantable cartilage constructs^11^. Gene expression changes in response to inflammatory cytokines (1 ng/mL IL-1α or 20 ng/ml TNF-*α*) were measured via qPCR (primers available in Supplementary Methods), and protein levels of secreted murine IL-1Ra were measured via ELISA (Duoset, R&D Systems).

### nanoCT Imaging of Bioartificial Implants

Implants were fixed overnight in 2.5% glutaraldehyde and 2% paraformaldehyde in 0.1 M cacodylate buffer with 2 mM CaCl_2_. Secondary fixation was performed using 1% OsO4 in 0.1 cacodylate buffer for 1 hour at room temperature. Excess fixative was washed using deionized water, and the samples were incubated in Lugol’s iodine for 72 hours at room temperature to further enhance contrast. Samples were embedded in 2% agarose and imaged on a Zeiss Xradia Versa 540 X-ray microscope using a 0.4x flat panel detector. Images were reconstructed and presented as shaded volume renderings.

### In Vivo Luciferase Assays

Ccl2-Luc constructs were implanted on the dorsal aspect of F1 K/BxN mice, which spontaneously develop chronic arthritis, to allow for longitudinal bioluminescence imaging of luciferase activity of the scaffolds *in vivo* daily for one week, and weekly thereafter over a 5-week period. C57BL/6 mice implanted with Ccl2-Luc scaffold constructs were challenged with K/BxN serum and imaged daily for 7 days (Supplementary Methods).

### Cell Viability Assays

Cell survival and apoptosis were assessed using the Live/Dead^®^ Cell Viability/ Cytotoxicity Kit for mammalian cells (Invitrogen/Molecular Probes, Carlsbad, California, USA). Live cells were labeled with calcein AM and dead cells were labeled with ethidium homodimer-1 bound to DNA. Labeled constructs were imaged using confocal microscopy (LSM 880, Zeiss, Thornwood, NY, USA)^11^.

### K/BxN Model of Inflammatory Arthritis

Male K/BxN transgenic mice and female B6.I-A^g7^ mice were intercrossed to generate F1 that spontaneously developed arthritis beginning at about 4 weeks of age and lasting for > 20 weeks^10^. To induce STA, C57BL/6 male and female mice received either dorsal subcutaneous Ccl2-IL1Ra scaffold implants (n=8 scaffolds, 8 animals, Experimental group), Ccl2-Luc scaffolds or no treatment (Control groups, n=8-13). One week after subcutaneous scaffold implantation, mice were challenged with 150-175µL of K/BxN serum delivered by retroorbital injection to induce arthritis^10,28,29^. Disease activity (clinical score and ankle thickness) and pain sensitivity (algometry and Electronic Von Frey) were assessed daily for 1 week. Mice were then sacrificed, and serum and hindpaws were collected for analysis. All procedures were approved by IACUC at Washington University in St. Louis.

### Histological Analysis

Paws were harvested on day 7 after serum transfer, fixed in 10% neutral buffered formalin for 48 h and stored in 70% ethanol, before decalcification in EDTA solution and processing for paraffin embedding. Sections (5 µm) were stained with Hematoxylin and Eosin (H&E) or toluidine blue. Inflammatory cells infiltrating the synovial lining and the joint cavity were enumerated in 8–10 random fields per section using H&E images acquired at 400x magnification. Proteoglycan loss in the cartilage was scored on toluidine blue-stained sections on a scale from 0–3, ranging from fully stained cartilage (score = 0) to fully destained cartilage (score = 3) as previously described^30^. Scoring was performed by an observer blinded to the treatment.

### Quantitation of IL-1Ra and Proinflammatory Mediators

Levels of IL-1Ra in serum and paws were assessed by ELISA (Quantikine-IL1Ra, R&D Systems). Levels of paw and serum inflammatory mediators were evaluated using Luminex® (Mouse High Sensitivity T-cell discovery array 18-Plex, Eve Technologies, Calgary, AB, Canada).

### MicroCT Analysis of Bone Erosion

To measure bone morphological changes, intact hind paws were scanned by microcomputed tomography (microCT, SkyScan 1176, Bruker) with a 9 µm isotropic voxel resolution at 50 kV, 500 µA, 980 ms integration time, 3 frame averaging, and 0.5 mm aluminum filter to reduce the effects of beam hardening. Images were reconstructed using NRecon software (with 20% beam hardening correction and 15 ring artifact correction). Hydroxyapatite calibration phantoms were used to calibrate bone density values (g/cm^3^). Parameters reported are: bone surface to bone volume (BS/BV), bone fraction (bone volume/total volume; BV/TV), and bone mineral density (BMD; g/cm^3^).

### Statistical Analysis

Sample size was determined based on a mean of 10 ± 2 for KRN control animals and 6 for IL-1Ra treated animals. Based on an alpha of 0.05 and 80% statistical power (1-β) a priori a sample size of 4 animals/treatment group was needed to observe this effect. Outcomes were evaluated by two-way Student’s t-test, one-way Mann-Whitney *U* test or one-, two-, three-way repeated measures, or three-way ANOVA with Tukey’s *post hoc* test or Sidak Correction to assess differences between groups, treatments, scaffold types, time, or a combination of those factors. Pearson or Spearman correlations were calculated between serum levels of IL-1Ra and outcomes (p ≤ 0.05). Investigators were not blinded to the clinical score or ankle thickness measurements. All other assessments and analyses were performed blinded.

## Acknowledgments

We thank Dr. Charles Gersbach for important discussions in the early stages of this work and Sara Oswald for providing technical writing support for the manuscript. This study was supported in part by grants NIH grants AR50245, AR48852, AG15768, AR48182, AG46927, OD10707, DK108742, AR073752, AR057235, AR067491, the Arthritis Foundation, and the Nancy Taylor Foundation for Chronic Diseases, NIH P50 CA094056 (Molecular Imaging Center) and NCI P30 CA091842 (Siteman Cancer Center Small Animal Cancer Imaging shared resource).

## Author Contributions

Y.R.C., K.H.C., C.T.N.P., and F.G. conceived the project, Y.R.C., K.H.C., C.L.W., J.M.B., C.T.N.P. and F.G. designed the experiments. Y.R.C., L.P., A.K.R., F.T.M., and C.L.W. conducted *in vitro* studies. *In vivo* experiments and data analyses were performed by Y.R.C, K.H.C., L.E.S., L.P., A.K.R., N.S.H., and C.L.W. Y.R.C., K.H.C., and F.G. wrote the manuscript. All the authors read and approved the final manuscript.

## Data Sharing

The data that support the findings of this study are available from the corresponding author upon reasonable request.

## Supplementary Figures

**Suppl. Fig. 1.**
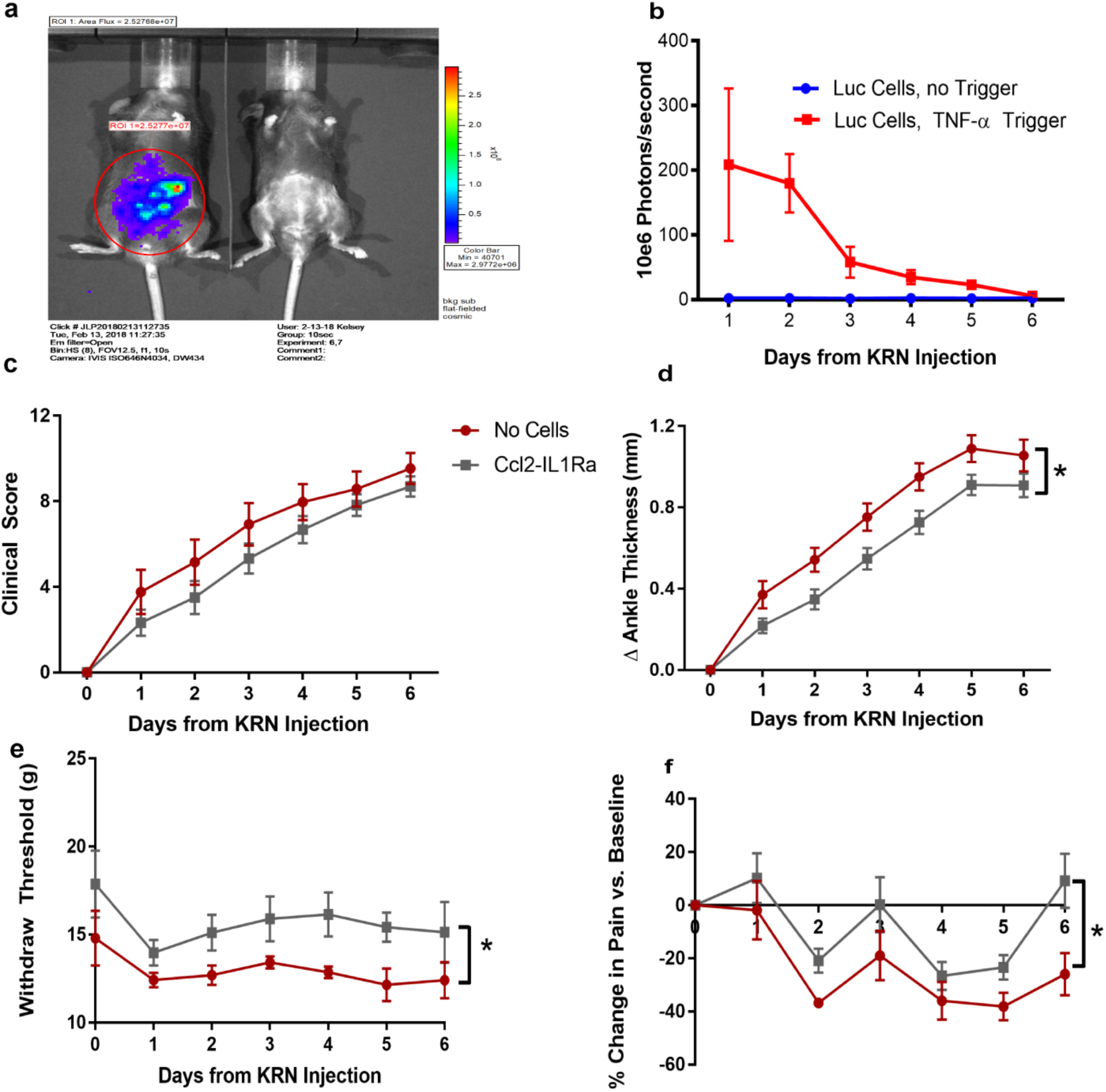
To assess an alternative cell delivery approach, Ccl2-Luc cells were delivered by IP injection in 200µL of sterile PBS to K/BxN STA mice. **a**. The injected cells demonstrated rapid luciferase expression (within 1h) in response to supraphysiological levels of pro-inflammatory cytokines (a, left TNF-*α* challenge with Ccl2-Luc cells; right no TNF-*α* challenge with Ccl2-Luc cells). **b**. Luciferase expression decreased over 6 days. **c**. With K/BxN serum transfer after IP injection of Ccl2-IL1Ra cells, significant mitigation of clinical scores was not observed (p=0.10), but **d**. did result in significant mitigation of ankle thickness with TNF-*α* challenge. IP injected Ccl2-IL1Ra cells also mitigated pain measured by **e**. algometry and **f**. Electronic Von Frey respectively in K/BxN STA mice, but to a lesser degree than when compared with the tissue-engineered Ccl2-IL1Ra cartilage scaffold constructs (e, f). All values represent mean ± SEM. *p < 0.05.

**Suppl. Fig. 2.**
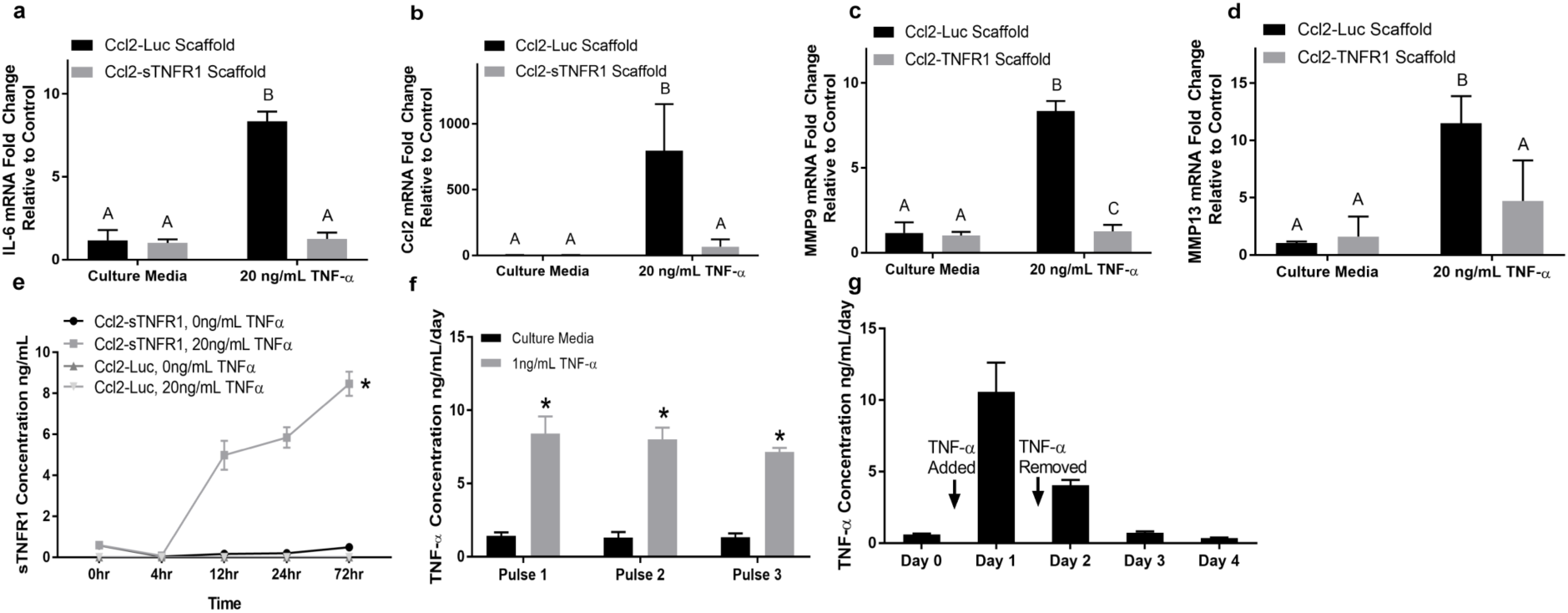
Ccl2-sTNFR1 anti-cytokine cartilaginous constructs mitigated mRNA levels for pro-inflammatory mediators **a**. *Il6*, **b**. *Ccl2*, **c**. *Mmp13*, and **d**. *Mmp9* when challenged with 20 ng/mL TNF-*α*. **e**. Ccl2-sTNFR1 constructs produced sTNFR1 in response to cytokine challenge. **f**. This response was repeatable over multiple cycles of stimulation. **g**. The levels of sTNFR1 were reduced to baseline levels after cytokine withdrawal. All values represent mean ± SEM. * and different letters indicate p<0.05.

**Suppl. Fig. 3.**
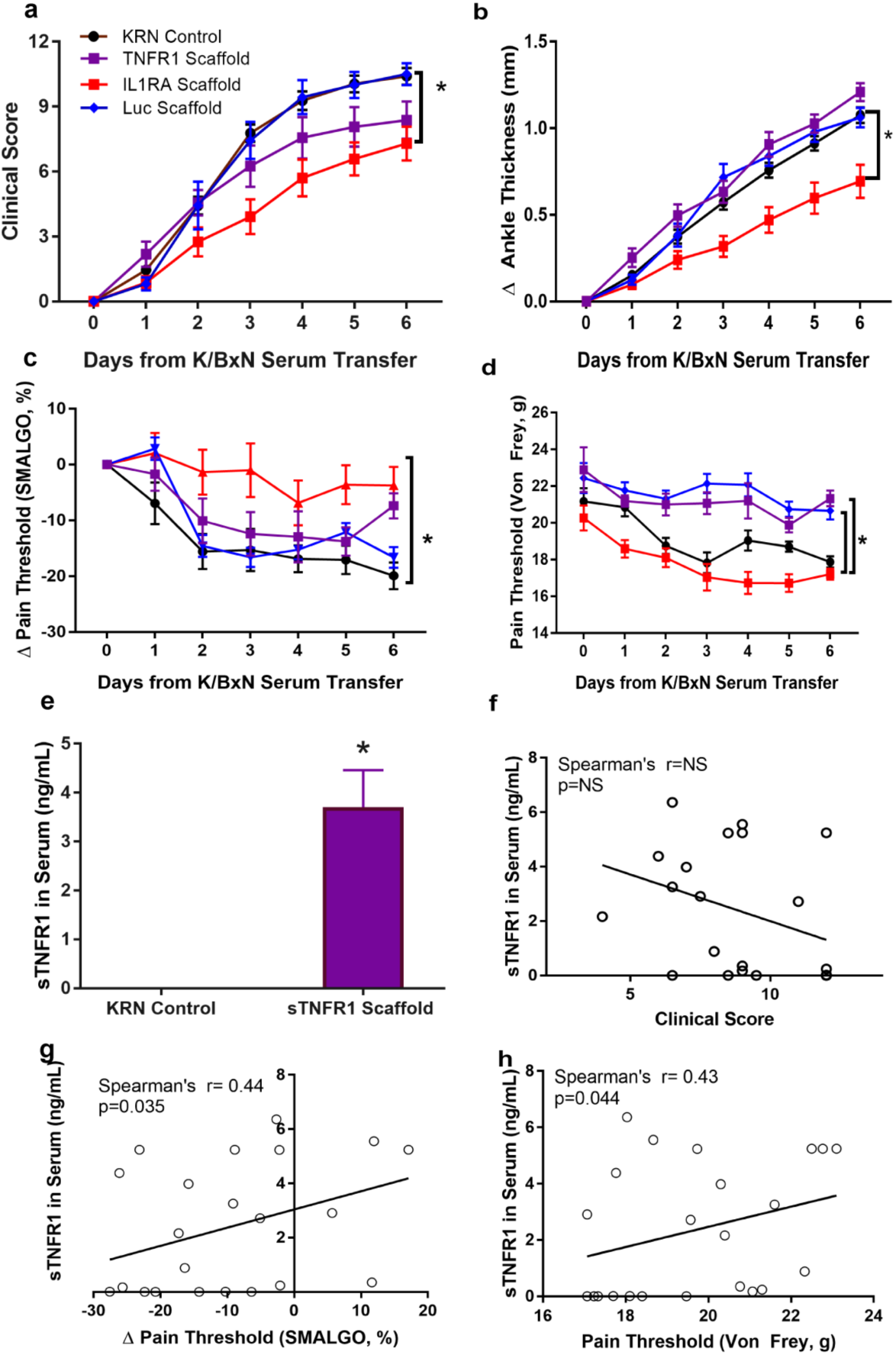
K/BxN serum transfer arthritis mice implanted with Ccl2-IL1Ra constructs demonstrated **a**. reduced clinical scores and **b**. ankle thickness when compared to mice receiving Ccl2-Luc constructs, Ccl2-sTNFR1 constructs, or mice receiving no implants. **c**. Mice with Ccl2-IL1Ra implants maintained their pain thresholds, indicated by algometry **d**. Both Ccl2-sTNFR1 and Ccl2-IL1Ra groups maintained pain thresholds using the Electronic Von Frey pain test. **e**. At sacrifice, significant increases in sTNFR1 concentration in serum from mice receiving Ccl2-sTNFR1 implants was observed compared to mice with no implants. **f**. No significant correlation was observed between sTNFR1 levels and the clinical score, but sTNFR1 concentration in serum were correlated with **g**. algometry and **h**. Von Frey pain thresholds. All values represent mean ± SEM. *p < 0.05.

**Suppl. Fig. 4.**
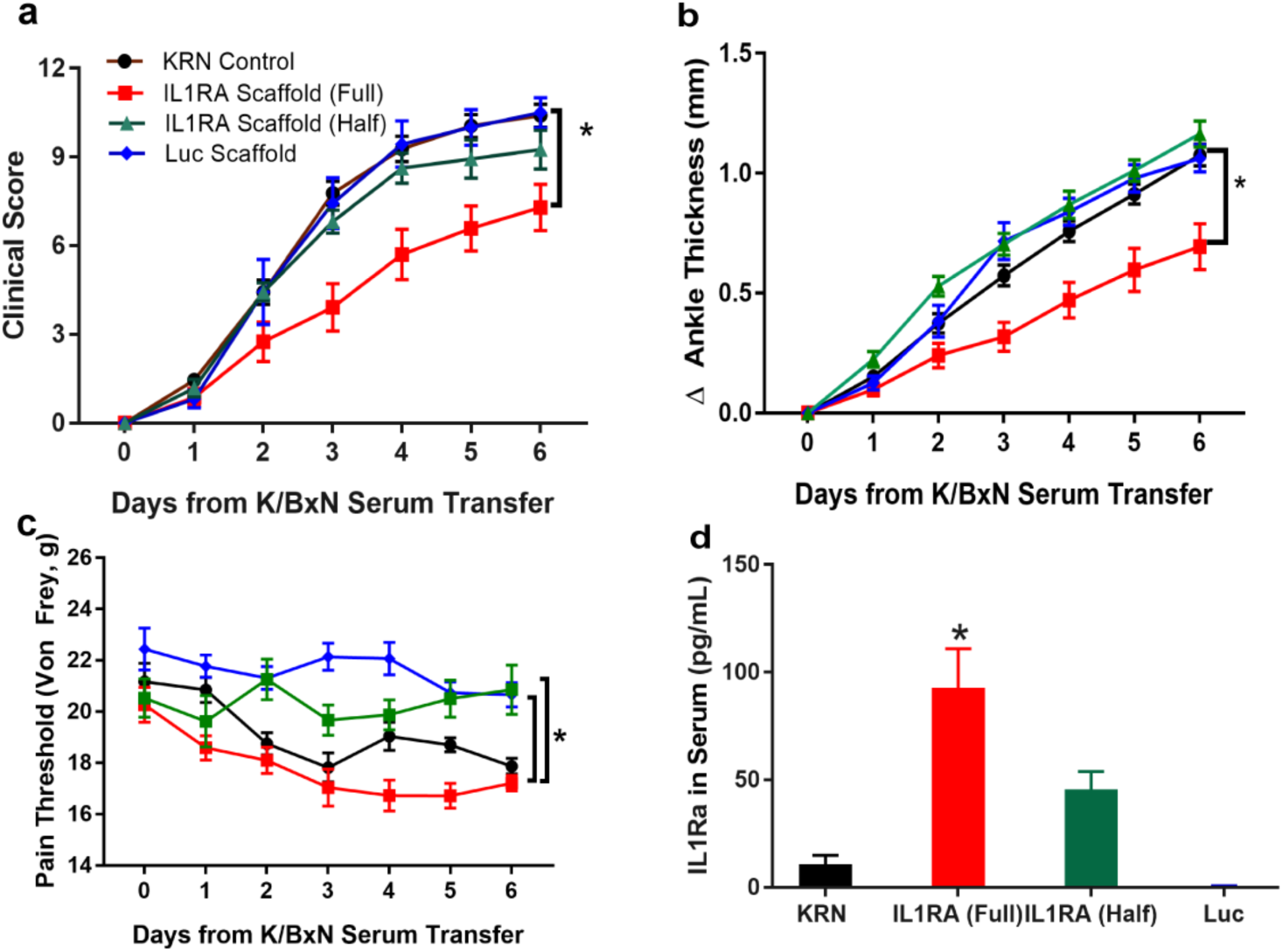
A dose-response was observed in genome-engineered constructs depending on the number of cells implanted (Ccl2-Luc, Ccl2-IL1Ra 8 implants [8e6 cells total, full dose] or Ccl2-IL1Ra 4 implants [4e6 total, half dose]) were implanted subcutaneously in mice with K/BxN serum transfer arthritis. **a**. Mice receiving Ccl2-IL1Ra implants demonstrated reduced clinical scores, and **b**. reduced ankle thickness as compared to mice receiving Ccl2-Luc implants, and both mice receiving half dose and no implants. **c**. Both Ccl2-IL1Ra treated groups maintained their pain thresholds, indicated by algometry. **d**. Approximately half the concentration of IL-1Ra was observed in the serum of the Ccl2-IL1Ra half dose group on average compared to Ccl2-IL1Ra full dose. Only the Ccl2-IL1Ra full dose group demonstrated significant increases in circulating IL-1Ra compared to mice with no implants. All values represent mean ± SEM. *p < 0.05.

## Supplementary Methods

### In Vivo Luciferase Assays

*In vivo* bioluminescence imaging was performed on the days indicated on an IVIS Lumina (PerkinElmer, Waltham, MA, USA; Living Image 4.2, 1 min exposure, bin 8, FOV 12.5cm, f/stop 1, open filter). Hair was removed from the area prior to imaging. Mice were injected intraperitoneally with D-luciferin (150 mg/kg in PBS; Gold Biotechnology, St. Louis, MO, USA) and imaged using isoflurane anesthesia (2% vaporized in O_2_). Total photon flux (photons/sec) was measured from fixed regions of interest (ROIs) over the lower backs of the mice where the scaffolds were located using Living Image 2.6.

### K/BxN Serum Transfer Experiments

The clinical score was assessed on a scale of 0–3 (0=no swelling or erythema, 1=slight swelling or erythema, 2=moderate erythema and swelling in multiple digits or entire paw, 3=pronounced erythema and swelling of entire paw; maximum total score of 12)^30^. The change from baseline in ankle thickness was determined daily by dial calipers, and an average change in the ankle thickness was determined for each mouse from the two hind paw measurements.

Animals were acclimatized to the behavioral testing room prior to assessment. Algometry was performed using a pressure-based analgesimeter (SMALGO, Bioseb, Vitrolles, France) by applying a progressive force over the ankle joint, and the stimulation was increased until a reaction (shudder or vocalization) from the animal was obtained. The maximum force was then automatically recorded and analyzed.

For Electronic Von Frey paw tests, mice were placed in acrylic cages (12 x 10 x 17 cm high) with a wire grid floor 15-30 min before testing in a quiet room. During this adaptation period, the hind paws were poked 2-3 times. Before hind paw stimulation, the animals were quiet, without exploratory movements or defecation, and not resting on their paws. In these experiments, we used a pressure-meter which consisted of a hand-held force transducer fitted with a 0.5 mm^2^ polypropylene tip (electronic von Frey anesthesiometer, IITC Inc., Life Science Instruments, Woodland Hills, CA, USA). Then, a tip was applied against the central edge of the animal hind paw and the intensity of the stimulus was automatically recorded when the paw was withdrawn. The stimulation of the paw was repeated until the animal presented two similar measurements.

## References

1 Gottenberg, J. E. et al. Non-TNF-Targeted Biologic vs a Second Anti-TNF Drug to Treat Rheumatoid Arthritis in Patients With Insufficient Response to a First Anti-TNF Drug: A Randomized Clinical Trial. JAMA 316, 1172–1180, doi: 10.1001/jama.2016.13512 (2016).

2 Tarp, S. et al. Risk of serious adverse effects of biological and targeted drugs in patients with rheumatoid arthritis: a systematic review meta-analysis. Rheumatology 56, 417–425, doi: 10.1093/rheumatology/kew442 (2017).

3 Brunger, J. M., Zutshi, A., Willard, V. P., Gersbach, C. A. & Guilak, F. Genome Engineering of Stem Cells for Autonomously Regulated, Closed-Loop Delivery of Biologic Drugs. Stem Cell Reports 8, 1202–1213, doi: 10.1016/j.stemcr.2017.03.022 (2017).

4 Joshi, N. et al. Towards an arthritis flare-responsive drug delivery system (vol 9, 2018). Nat Commun 9, doi: 10.1038/s41467-018-04346-x (2018).

5 Ye, H. et al. Self-adjusting synthetic gene circuit for correcting insulin resistance. Nat Biomed Eng 1, 0005, doi: 10.1038/s41551-016-0005 (2017).

6 Doudna, J. A. & Charpentier, E. Genome editing. The new frontier of genome engineering with CRISPR-Cas9. Science 346, 1258096, doi: 10.1126/science.1258096 (2014).

7 Gaj, T., Gersbach, C. A. & Barbas, C. F., 3rd. ZFN, TALEN, and CRISPR/Cas-based methods for genome engineering. Trends Biotechnol 31, 397–405, doi: 10.1016/j.tibtech.2013.04.004 (2013).

8 Hao, S. & Baltimore, D. The stability of mRNA influences the temporal order of the induction of genes encoding inflammatory molecules. Nat Immunol 10, 281–288, doi: 10.1038/ni.1699 (2009).

9 Murphy, M. B., Moncivais, K. & Caplan, A. I. Mesenchymal stem cells: environmentally responsive therapeutics for regenerative medicine. Exp Mol Med 45, e54, doi: 10.1038/emm.2013.94 (2013).

10 Kyburz, D. & Corr, M. The KRN mouse model of inflammatory arthritis. Springer Semin Immunopathol 25, 79–90, doi: 10.1007/s00281-003-0131-5 (2003).

11 Moutos, F. T. et al. Anatomically shaped tissue-engineered cartilage with tunable and inducible anticytokine delivery for biological joint resurfacing. Proc Natl Acad Sci U S A 113, E4513–4522, doi: 10.1073/pnas.1601639113 (2016).

12 Diekman, B. O. et al. Cartilage tissue engineering using differentiated and purified induced pluripotent stem cells. Proc Natl Acad Sci U S A 109, 19172–19177, doi: 10.1073/pnas.1210422109 (2012).

13 Atala, A. et al. Injectable alginate seeded with chondrocytes as a potential treatment for vesicoureteral reflux. J Urol 150, 745–747 (1993).

14 Makris, E. A., Responte, D. J., Paschos, N. K., Hu, J. C. & Athanasiou, K. A. Developing functional musculoskeletal tissues through hypoxia and lysyl oxidase-induced collagen cross-linking. Proc Natl Acad Sci U S A 111, E4832–4841, doi: 10.1073/pnas.1414271111 (2014).

15 Mauck, R. L., Wang, C. C., Oswald, E. S., Ateshian, G. A. & Hung, C. T. The role of cell seeding density and nutrient supply for articular cartilage tissue engineering with deformational loading. Osteoarthritis Cartilage 11, 879–890 (2003).

16 Vunjak-Novakovic, G. et al. Bioreactor cultivation conditions modulate the composition and mechanical properties of tissue-engineered cartilage. J Orthop Res 17, 130–138, doi: 10.1002/jor.1100170119 (1999).

17 Christensen, A. D., Haase, C., Cook, A. D. & Hamilton, J. A. K/BxN Serum-Transfer Arthritis as a Model for Human Inflammatory Arthritis. Front Immunol 7, 213, doi: 10.3389/fimmu.2016.00213 (2016).

18 Schett, G. & Gravallese, E. Bone erosion in rheumatoid arthritis: mechanisms, diagnosis and treatment. Nat Rev Rheumatol 8, 656–664, doi: 10.1038/nrrheum.2012.153 (2012).

19 Aimetti, A. A., Tibbitt, M. W. & Anseth, K. S. Human neutrophil elastase responsive delivery from poly(ethylene glycol) hydrogels. Biomacromolecules 10, 1484–1489, doi: 10.1021/bm9000926 (2009).

20 Brudno, Y. & Mooney, D. J. On-demand drug delivery from local depots. J Control Release 219, 8–17, doi: 10.1016/j.jconrel.2015.09.011 (2015).

21 Ehrbar, M. et al. Biomolecular hydrogels formed and degraded via site-specific enzymatic reactions. Biomacromolecules 8, 3000–3007, doi: 10.1021/bm070228f (2007).

22 Langer, R. Implantable controlled release systems. Pharmacol Ther 21, 35–51 (1983).

23 Brunger, J. M., Zutshi, A., Willard, V. P., Gersbach, C. A. & Guilak, F. CRISPR/Cas9 Editing of Murine Induced Pluripotent Stem Cells for Engineering Inflammation-Resistant Tissues. Arthritis Rheumatol 69, 1111–1121, doi: 10.1002/art.39982 (2017).

24 Farhang, N. et al. CRISPR-Based Epigenome Editing of Cytokine Receptors for the Promotion of Cell Survival and Tissue Deposition in Inflammatory Environments. Tissue Eng Part A 23, 738–749, doi: 10.1089/ten.TEA.2016.0441 (2017).

25 Deans, T. L., Grainger, D. W. & Fussenegger, M. Synthetic Biology: Innovative approaches for pharmaceutics and drug delivery. Adv Drug Deliv Rev 105, 1–2, doi: 10.1016/j.addr.2016.08.013 (2016).

26 Chen, Z., Bozec, A., Ramming, A. & Schett, G. Anti-inflammatory and immune-regulatory cytokines in rheumatoid arthritis. Nat Rev Rheumatol 15, 9–17, doi: 10.1038/s41584-018-0109-2 (2019).

27 Johnstone, B., Hering, T. M., Caplan, A. I., Goldberg, V. M. & Yoo, J. U. In vitro chondrogenesis of bone marrow-derived mesenchymal progenitor cells. Exp Cell Res 238, 265–272, doi: 10.1006/excr.1997.3858 (1998).

28 Korganow, A. S. et al. From systemic T cell self-reactivity to organ-specific autoimmune disease via immunoglobulins. Immunity 10, 451–461 (1999).

29 Kouskoff, V. et al. Organ-specific disease provoked by systemic autoimmunity. Cell 87, 811–822 (1996).

30 Zhou, H. F., Chan, H. W., Wickline, S. A., Lanza, G. M. & Pham, C. T. Alphavbeta3-targeted nanotherapy suppresses inflammatory arthritis in mice. FASEB J 23, 2978–2985, doi: 10.1096/fj.09-129874 (2009).

